# Adapt-A-Maze: An Open Source Adaptable and Automated Rodent Behavior Maze System

**DOI:** 10.1101/2021.06.05.447225

**Authors:** Blake S. Porter, Jacob M. Olson, Christopher A. Leppla, Éléonore Duvelle, John H. Bladon, Matthijs AA. van der Meer, Shantanu P. Jadhav

**Affiliations:** Neuroscience Program, Department of Psychology, and Volen National Center for Complex Systems, Brandeis University, Waltham, MA, USA; Department of Psychological and Brain Sciences, Dartmouth College, Hanover, NH, USA

## Abstract

Mazes are a fundamental and widespread tool in behavior and systems neuroscience research in rodents, especially in spatial navigation and spatial memory investigations in freely behaving animals. However, their form and inflexibility often restrict potential experimental paradigms that involve multiple or adaptive maze designs. Unique layouts often lead to elevated costs, whether financially or in terms of time investment from scientists. To alleviate these issues, we have developed an automated, modular maze system that is flexible and scalable. This open source Adapt-A-Maze (AAM) system will allow for experiments with multiple track configurations in rapid succession. Additionally, the flexibility can expedite prototyping of behavioral paradigms. Automation ensures less variability in experimental parameters and higher throughput. Finally, the standardized componentry enhances experimental repeatability within labs and replicability across labs. Our maze was successfully used across labs, in multiple experimental designs, with and without extracellular or optical recordings, in rats. The AAM system presents multiple advantages over current maze options and can facilitate novel behavior and systems neuroscience research.

**Significance statement:** We have developed an open source, modular maze system (the Adapt-A-Maze, AAM) that enables any lab interested in rodent behavior and cognition to create standard and unique mazes for their research. The AAM uses modular track pieces that can be combined to create a wide array of maze types. The AAM system also included reward wells with lick detection and pneumatic barriers. All aspects of the maze can be controlled via TTL signals to automate tasks and improve repeatability of experiments. The AAM system is expected to help labs quickly and inexpensively set up recordable experiments to advance our understanding of the neurophysiology underlying behavior and cognition.

## Introduction

Rodents navigating mazes is a longstanding and widespread method for investigating behavior, cognitive processes, and neurophysiology (for review, see Wijnen et al., 2024). Mazes leverage the fact that navigating to rewards or away from danger is a natural task for both rats and mice. Beginning over a century ago at the advent of the 20^th^ century (Small, 1901), maze experiments have resulted in landmark discoveries regarding cognitive processes of learning & memory underlying navigation (Tolman et al., 1946) and its neural substrates (O’Keefe & Dostrovsky, 1971; Olton & Samuelson, 1976). Currently, standard maze designs are pervasive and underlie foundational tasks in learning and memory, decision-making, and even anxiety (Barnes, 1979; Frank et al., 2000; Handley & Mithani, 1984; Morris et al., 1982; O’Keefe & Dostrovsky, 1971; Olton & Samuelson, 1976). In addition, unique mazes are constantly being designed to test specific questions in these fields (Ainge et al., 2007; Böhm & Lee, 2020; Knierim et al., 2000; Nitz, 2006; Olson et al., 2017; Porter et al., 2018; Steiner & Redish, 2014; Tanila et al., 2018; Wijnen et al., 2024; Wilson et al., 2015). Looking forward, the ability to rapidly switch environments while monitoring neural activity continuously is a growing desire in systems neuroscience to investigate contextual memory and representations, spatial remapping, and flexible decision making (Awh et al., 2024; Huelin Gorriz et al., 2023; Karlsson & Frank, 2009). The explosion of neural data recording capabilities is supported by the advances in machine learning techniques for analysis, giving increasing value to complex behavioral datasets.

Current maze designs limit the ability for multiple maze environment recordings as part of one experiment. Simple mazes often consist of one piece, with limited or no flexibility for other maze configurations. If multiple maze configurations are desired, they must all be made, stored, and then moved into and out of place in the room. This limits potential experiments and slows research progress. Alternatively, scientists will spend months engineering complex, automated mazes to address a specific question. Despite the time invested, these mazes are not often amenable to flexibly changing into other maze configurations, and so the same cost in design and manufacturing must be incurred again for the next experiment.

To address these issues, we have developed the Adapt-A-Maze (AAM), an open-source automated maze system that uses standardized track pieces to create flexible and scalable maze environments for rodents. The maze can be rapidly adapted to facilitate experiments that use multiple configurations, switching between shapes on the scale of minutes. Standardized reward wells are integrated into the track pieces and support automated lick detection and reward delivery along with quick, effective cleanup. Automated movable barriers can be placed between any track components, allowing additional environmental manipulations even during active behavior. Careful consideration of component selection enables artifact free electrophysiological recordings. Being open source, labs across the world could more easily replicate the experiments of other labs using the AAM. The AAM will allow researchers to complete better experiments faster, with greater repeatability, and at lower cost, supporting the advance of our understanding of rodent behavior and neurophysiology.

## Materials and Methods

### Materials Availability

All resources are made available on our Gitlab repository: https://gitlab.com/jadhavlab/AdaptAMaze/-/tree/main?ref_type=heads. This includes schematics, 3D printer design files and print files, track piece sheet metal SolidWorks files, PCB files, SpikeGadgets ECU scripts for automating simple tasks, and a detailed parts list. We’ve endeavored to use, or at least publish, file types that can be opened and modified in free software. Our goal is to make it so anyone can clone our repository and get their own AAM running in a few weeks. We look forward to users making modifications, creating new track pieces, and developing additional components to add to the repository.

### Code Accessibility

We have provided some example Spikegadgets Statescript programs can be found in the repository (https://gitlab.com/jadhavlab/AdaptAMaze/-/tree/main/Example%20Statescript%20programs?ref_type=heads).

### AAM Methods

Our parts list has detailed ordering and fabrication notes on applicable parts. **Figures 2** and **2-1** show the schematics of a full system. **Figures 3-1, 3-2**, and **4-1** show detailed pictures of all components and their part IDs. Part IDs and Excel sheets are color coded by the system they belong to.

**Figure 1:**
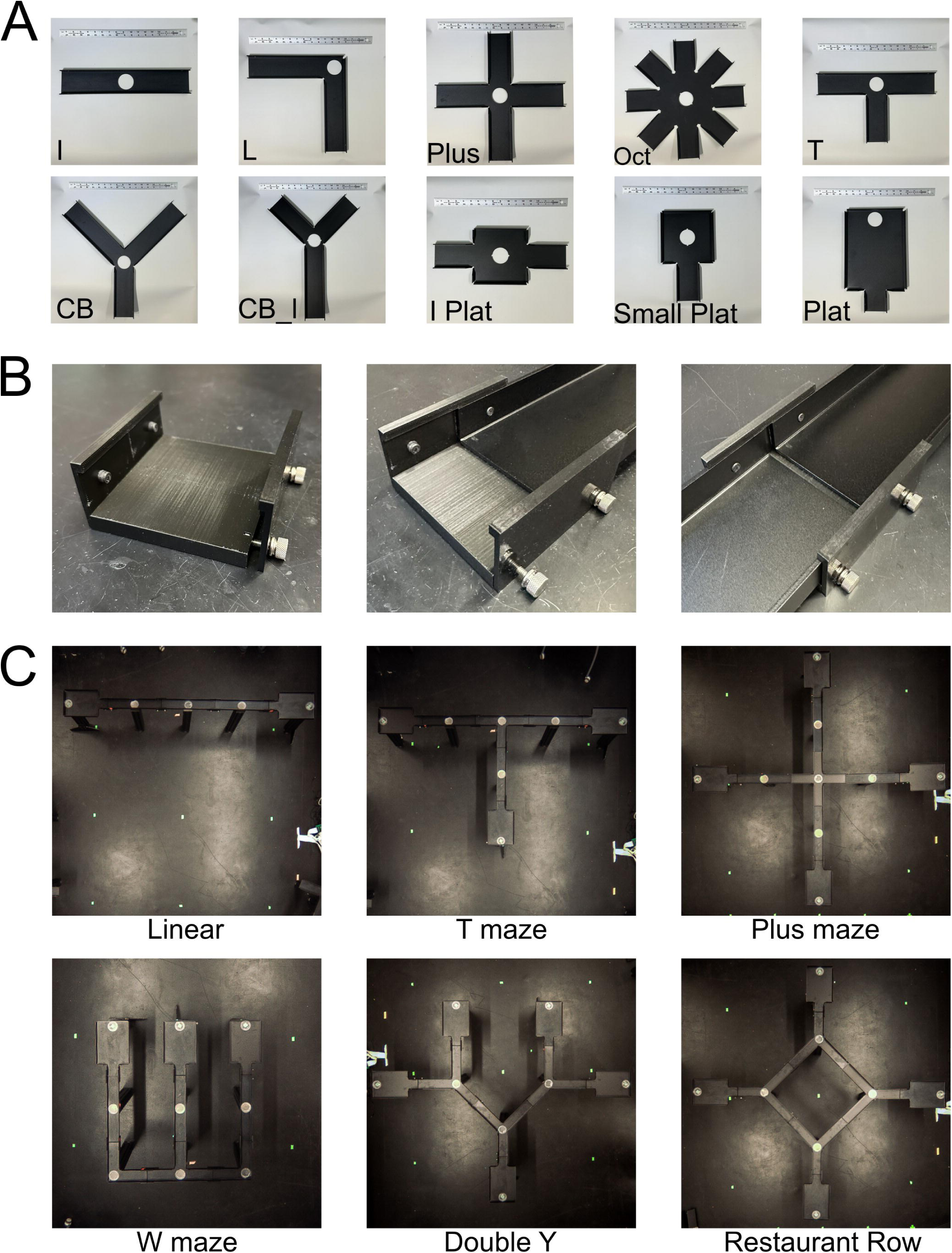
Adaptable Maze System. **A)** Mazes are constructed by combining track pieces into the desired shapes. Pictured are the 10 most common track piece shapes. Depicted ruler is 18 inches. **B)** Track joints support and join track pieces. **C)** Six example mazes, including common mazes from the neurophysiology literature. Only track pieces are shown; see **Figure 1-1** for fully assembled complex mazes. *CB = Corner Branch*, *Oct = Octagon, Plat = Platform*.

**Figure 1-1:**
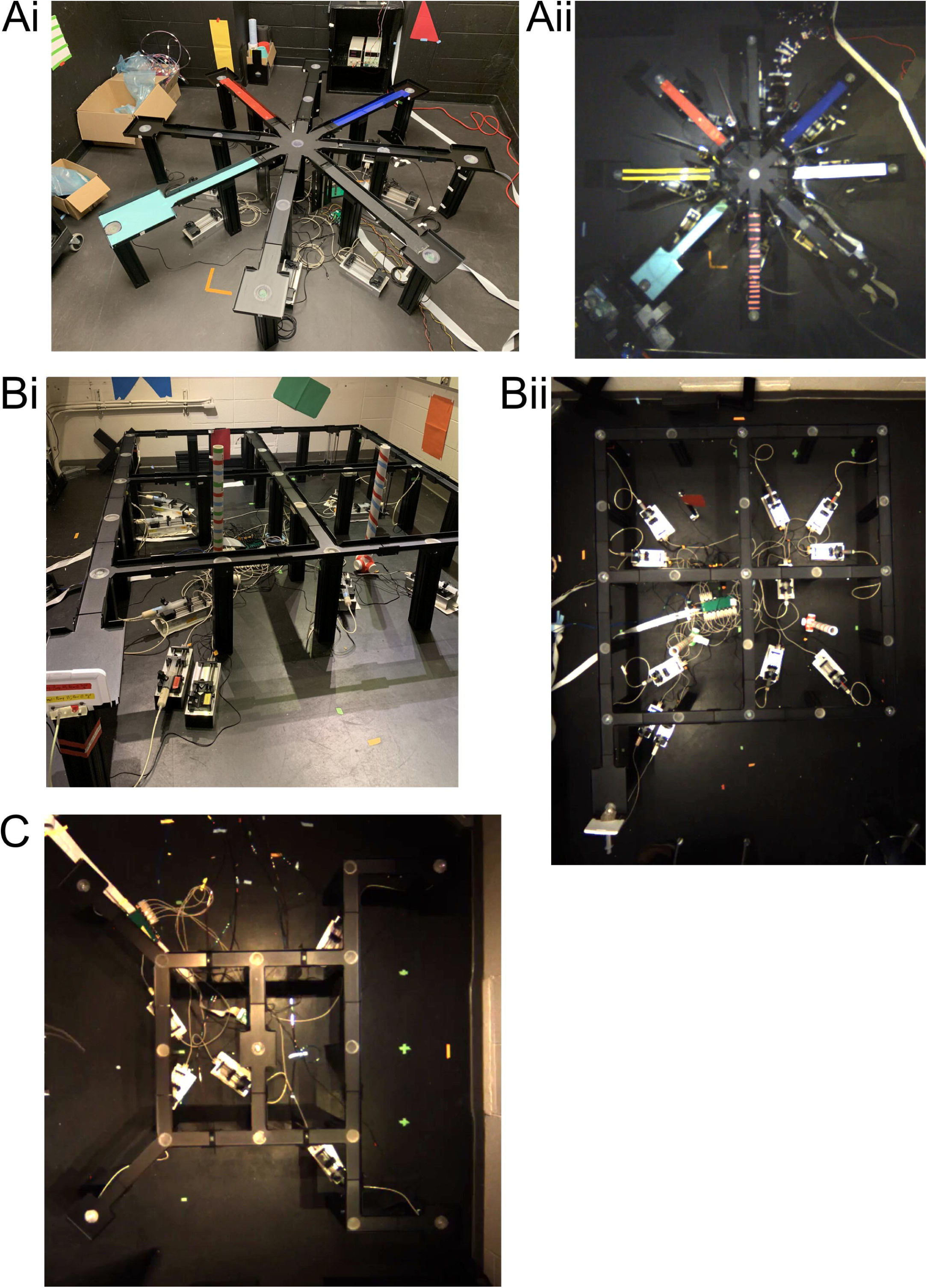
Complex maze setups. Members of the lab have implemented a variety of complex mazes in addition to common configurations (e.g. linear, W, T; Figure 1C). **A)** Classic radial arm maze used for a transitive inference task. Reward wells were located at the end of each arm. Barriers were placed at the start of each arm to control access to arms. Various materials were attached to the track pieces of each arm for unique local cues. **B)** Large (90” x 108”) 5×5 grid maze with a home arm (bottom left **Bi & Bii**) used for a memory schema paradigm. Reward wells were on each “node” of the grid. **C)** Large, complex maze with rewards available at a home location and four outer arms. Multiple routes could be taken from home to the outer arms. Available routes were manipulated via barriers and/or moving the home arm between sessions to test spatial memory.

**Figure 2:**
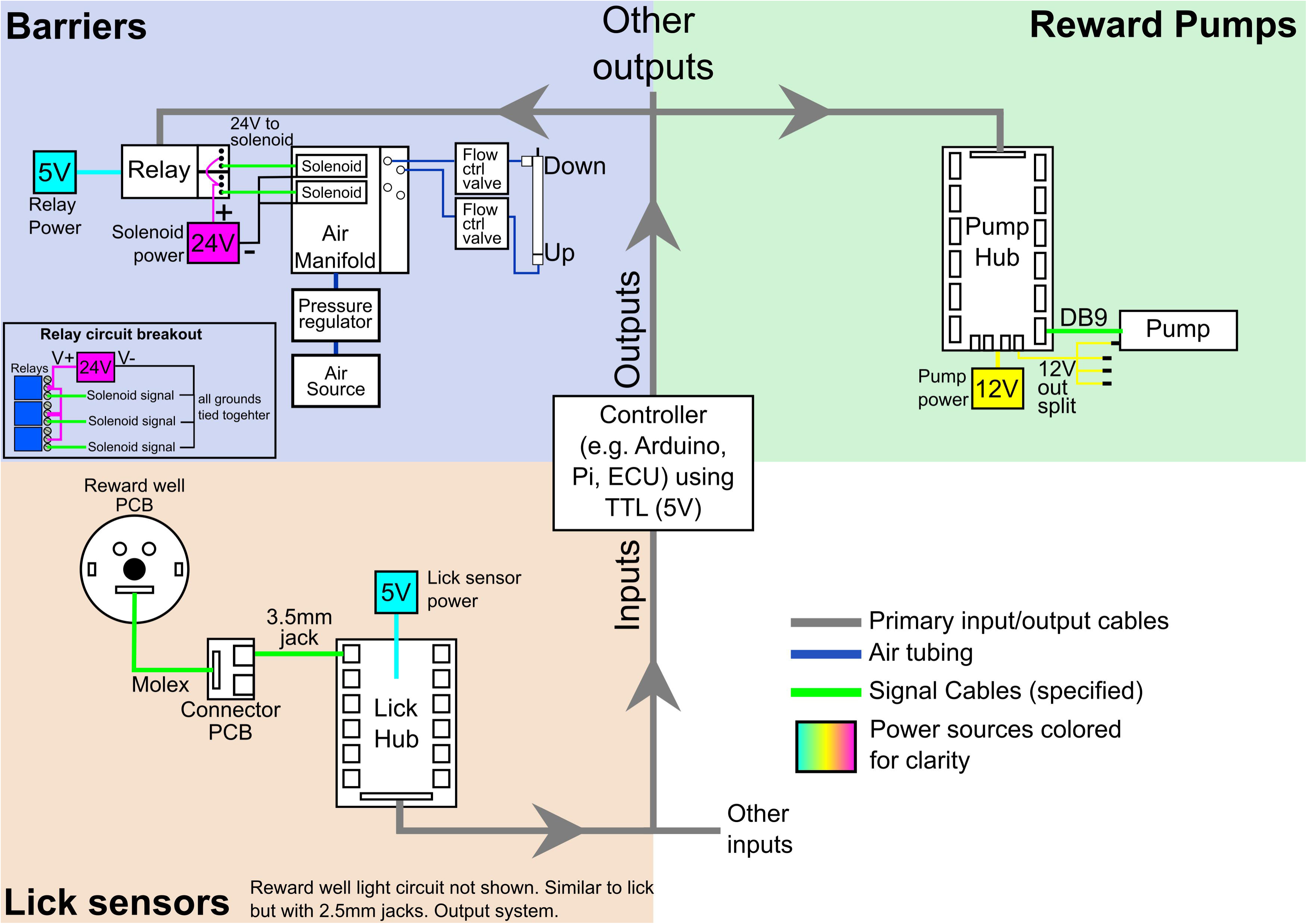
Simple schematic of an example AAM DIO setup. The depicted setup contains reward pumps, lick sensors, and barriers. A similar schematic but with pictures can be seen in **Figure 2-1**.

**Figure 2-1:**
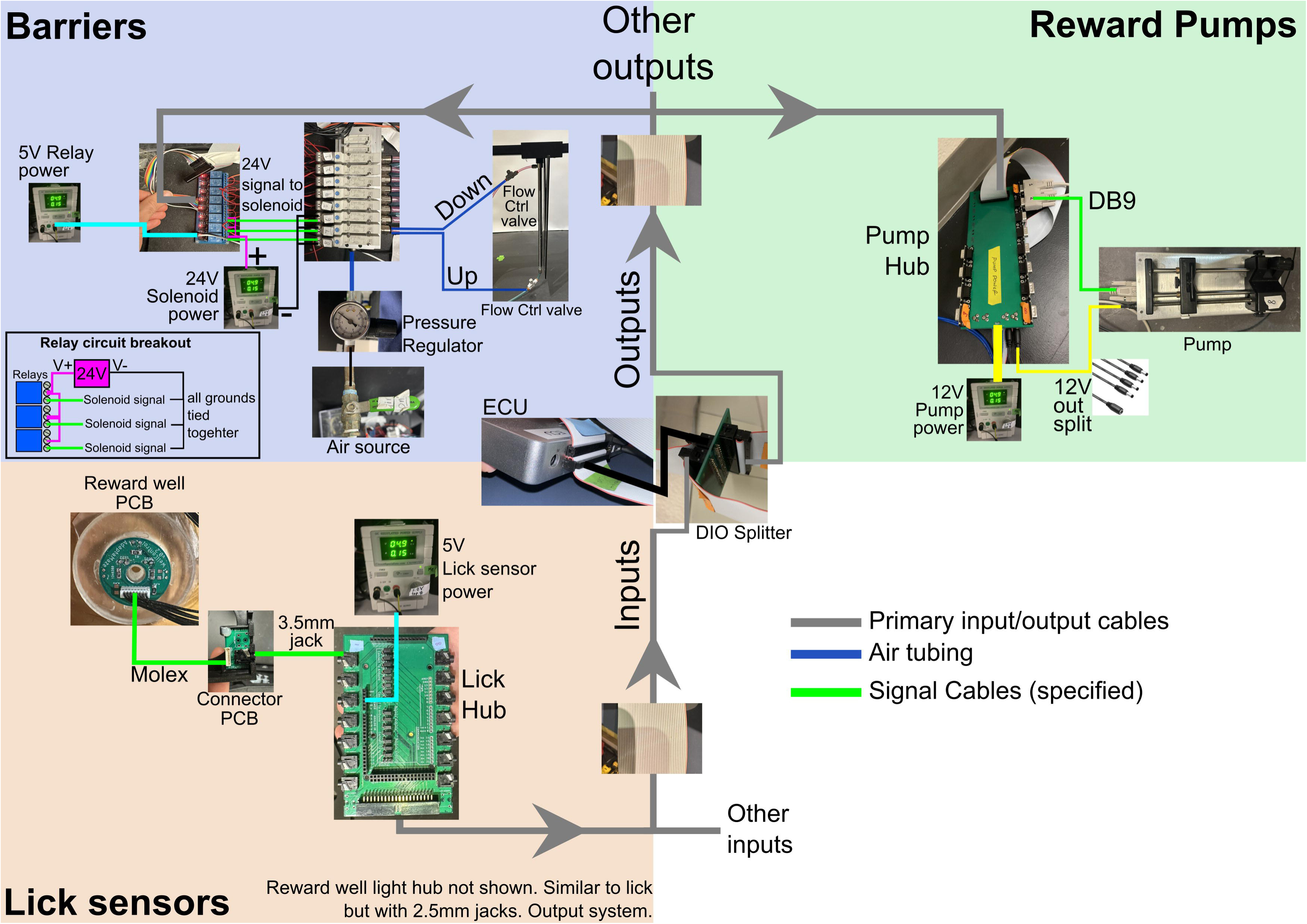
Picture-based schematic of AAM DIO system. Same as Figure 2 using SpikeGadget’s ECU as the controller.

**Figure 3:**
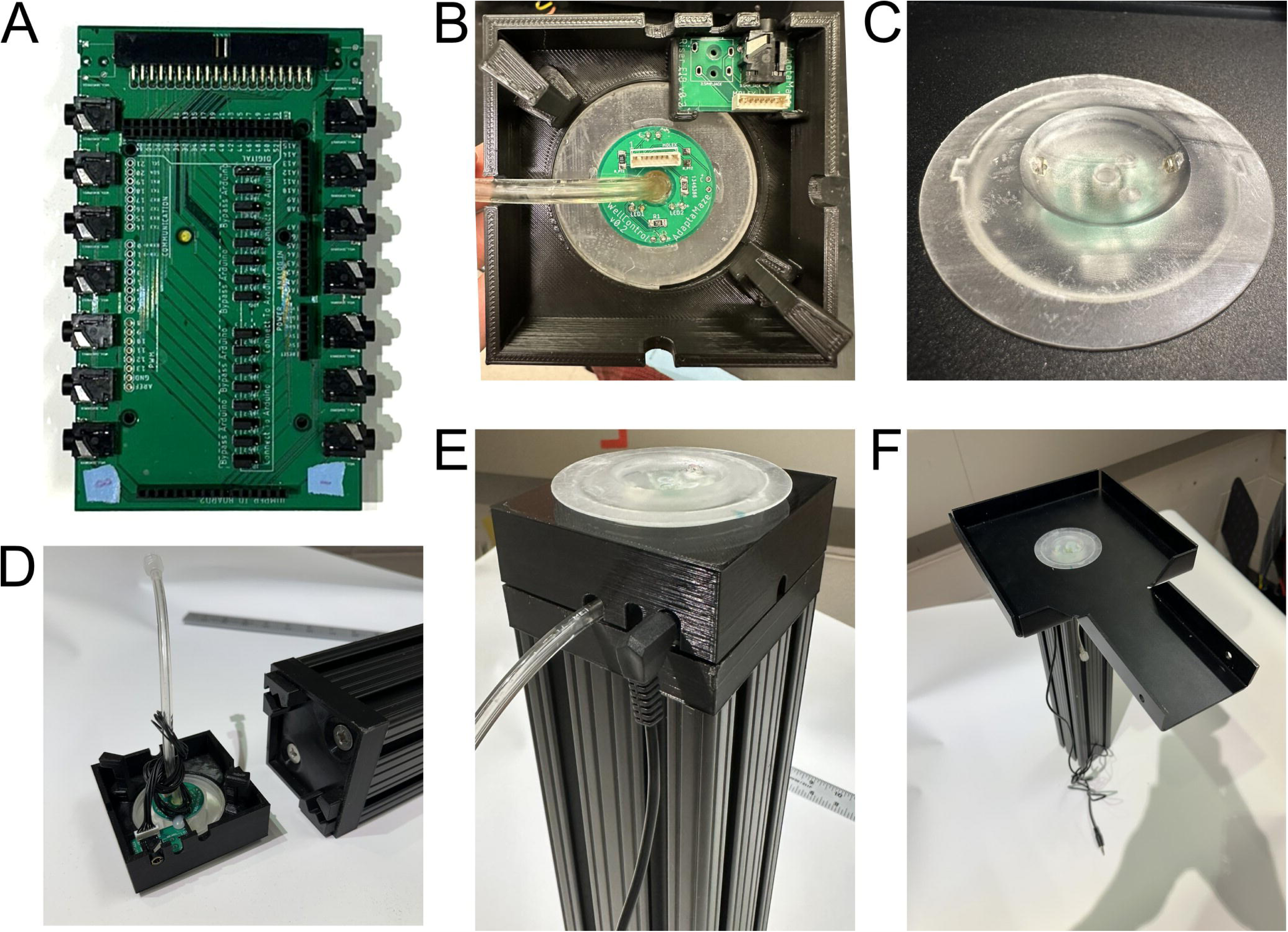
Integrated Automated Reward Wells. **A)** Reward well detection command PCB to route lick sensor cables from reward wells (shown in B) to ECU/controller and handle power to LEDs. **B)** Underside of constructed reward well and leg connector. No Wires shown. **C)** Example reward well with infrared (IR) emitter and phototransistor aligned across the liquid spout. **D)** Assembled reward well in connector (as in B) and leg with baseplate. **E)** Constructed “leg stack” with no track piece. **F)** Fully assembled modular track piece. Detailed parts breakouts for the reward well, track joint, and reward pumps are shown in **Figures 3-1 and 3-2**.

**Figure 3-1:**
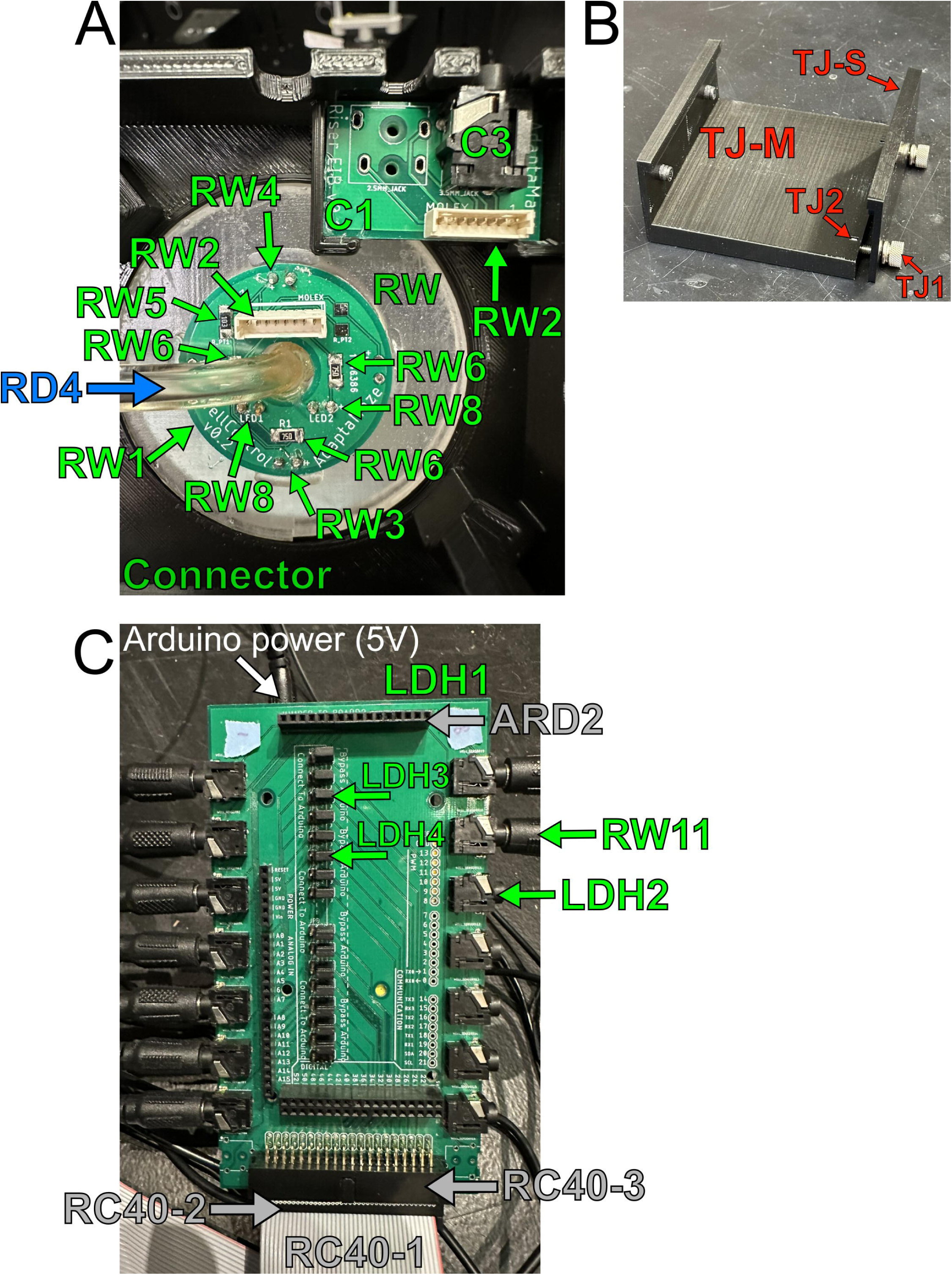
Reward well and track joint parts. Part breakouts for **A)** reward well & connector, **B)** track joint, and **C)** lick detection hub seen in Figure 3. Part IDs reference the parts list. Part ID colors correspond to the primary system they are associated with and are consistent with the Parts List’s tab colors. Red = track parts, green = reward wells, blue = reward delivery, grey = DIO connections.

**Figure 3-2:**
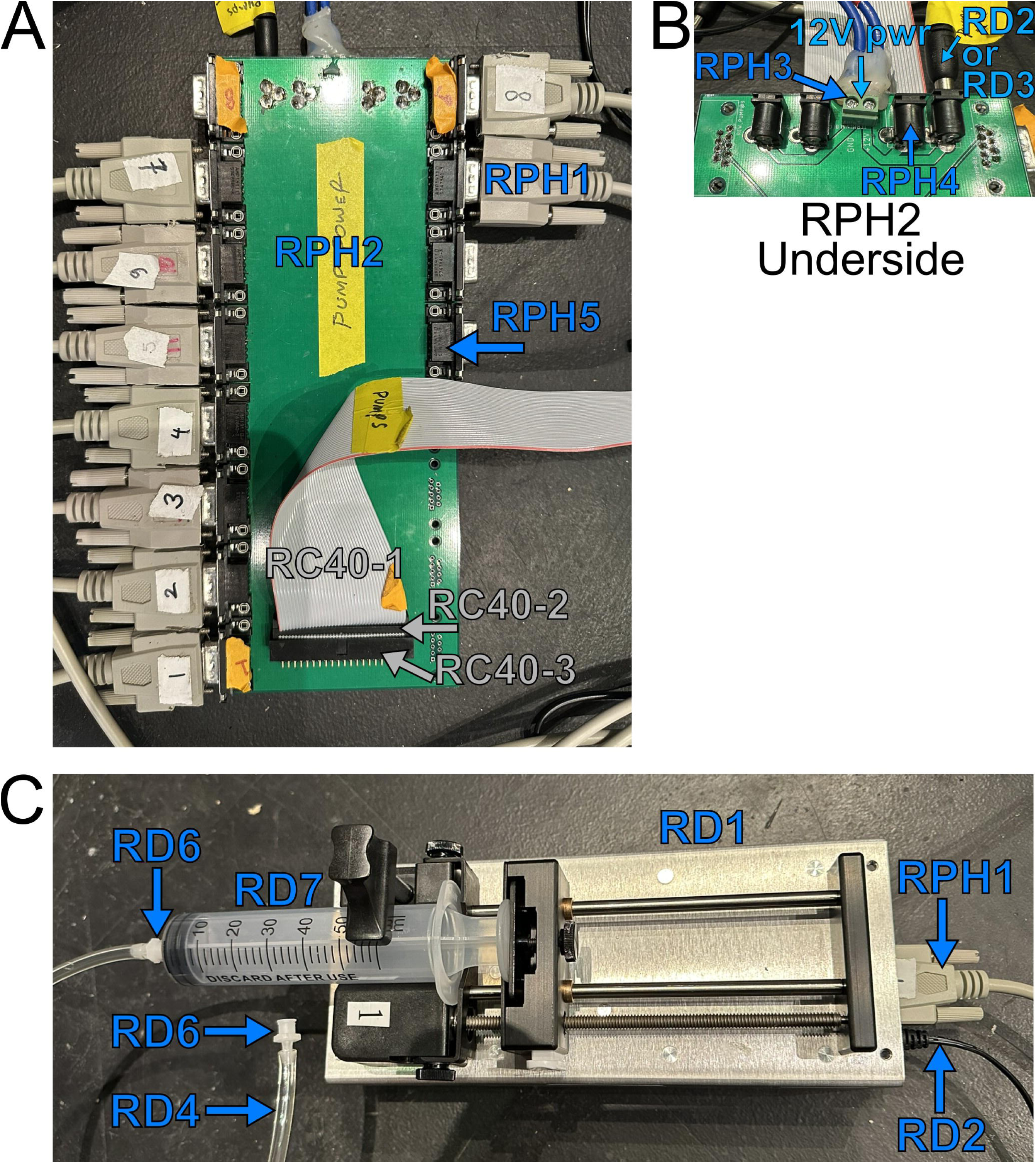
Reward pump parts. Part breakouts for **A)** Reward Pump hub, **B)** underside of reward pump hub, and **C)** syringe pump for Reward system (Figure 3). Part IDs reference the parts list. Part ID colors correspond to the primary system they are associated with and are consistent with the Parts List’s tab colors. Red = track parts, green = reward wells, blue = reward delivery, grey = DIO connections.

#### Track Pieces

Animals move on an interlocking system of custom anodized aluminum track pieces to form the maze environment. Track pieces are all 3” (7.6 cm) wide with 7/8 in (2.2 cm) walls and come in variable shapes and lengths primarily designed for an 18 in grid (**Figure 1**). Track pieces are connected using custom 3D-printed (Ender3 Pro) polylactic acid (PLA) plastic track joints with 0.25 in (0.6 cm) gaps between tracks (**Figure 1B**). Barriers necessitate wider gaps of 3 in. Barrier track joints can be used throughout to maintain grid spacing. Each track piece has a 2.5 in (6.4 cm) hole for insertion of a custom reward well or plug (**Figure 3A**) manufactured in-house using stereolithography (SLA) 3D-printing (Form2 and Form3 printers with clear resin, FormLabs Inc.).

#### Leg assembly

Each individual track piece is supported by its own leg assembly (**Figure 3F**) using a custom quick-lock system (**Figure 3B**). Contained within the leg assembly is the necessary electronics for reward wells.

Legs are made from 3 in x 3 in T-slotted beam (part# 3030, 8020 Inc.) and a custom 3-D printed base plate (**Figure 3C**). Fifteen-inch-tall legs have been extensively used with rats with few adverse events (e.g., rats attempting to climb down). We have utilized legs as tall as 36 in to further discourage climbing down. Legs of any height can be ordered to suit experimental needs. The base plate is attached to the top of the T-slotted beam to lock in with the connector.

The leg assembly is constructed by inserting a reward well or plug through a track piece and screwing it onto the connector (**Figure 3D**). The track assembly can then be installed onto the leg using the quick-lock system, which requires insertion of the connector hooks into the base plate, a 10-degree twist into position, and a press down to lock onto the base plate.

#### Automated Reward System

An infrared beam break is integrated into the reward well to detect licks (**Figure 3C, 6E**), and reward delivery tubing is routed through the bottom of the well to deliver liquid reward. Port entry detection, licks, and reward delivery are automated using custom hardware and software connected into the SpikeGadgets environmental control unit. This setup allows for precise control and automation of custom experimental setups using high level programming languages such as python or MATLAB (Mathworks Inc.). However, other controllers capable of TTL (e.g. Arduino, Raspberry Pi, Neuralynx Inc. Digital Lynx, OpenEphys) could easily be used instead of SpikeGadgets.

In almost all of our tasks, we have implemented a lick requirement before a reward is dispensed. With the exception of initial training on a linear track, we require rats lick at a reward well for a certain number of licks before a reward is released. This requirement is a behavioral readout that the rat chose an option, rather than just travelled to an area with no intention of collecting a reward. Furthermore, the requirement protects against random footfalls or tails from triggering a reward. We generally we count beam breaks that last more than 25 ms but less than 250 ms as “real licks”. Occasionally we need to adjust these parameters, especially the top end duration, for specific animals. We have provided out shaping code to train the animals to lick on the Gitlab repository.

The current reward well designs do not have a drainage system. We recommend the above lick requirements to avoid erroneously dispensed rewards. However, there may be situations where rats do not consume all the dispensed rewards. If such a scenario is anticipated, it may be beneficial to track the reward wells that had reward dispensed but a consummatory bout of licking did not follow. This may lock out that well from dispensing more reward on the next visit. A drainage system may be possible but would require the modification of the reward well and likely the reward well PCB. Furthermore, while we have not tested this ourselves, the syringe pumps are capable of withdrawing which could remove unconsumed liquid.

#### Automated Barriers

Automated barriers can be integrated into the maze environment between any two track pieces for within-experiment adaptation of the environment (**Figure 4**). Barriers are integrated into the track joint pieces and rise from between the two pieces using a pneumatic cylinder. Air input to the cylinders is controlled via 24V solenoids connected to a relay board connected to the ECU (**Figure 2**). Any microcontroller capable of TTL could control the barriers’ states as well. While pneumatic cylinders are audibly loud, rats habituate to the sound within 1-2 exposure sessions (see **Movie 1**). Pneumatic cylinders avoid electrical interference from alternative methods such as stepper motors (which we tested to great lengths but found them to be unsatisfactory). Each barrier’s speed can be fine-tuned via flow control valves. At 12 PSI, we set our flow control valves so that the barriers move both up and down in 0.4 seconds. However, the air cylinders are rated for 250 PSI so the barriers’ speed could be made incredibly fast. The flow control valves could also be adjusted to move the barriers very slowly if needed (e.g., < 1 cm / s). If the experiment allows, we recommend only moving barriers when rats are at known safe location, such as licking at distant reward wells, to eliminate the possibility of injuring the animal. However, with careful barrier design and care to minimize any gaps in the track pieces, the moving barriers pose little risk to rats’ tails (see **Movie 1**). The barrier system only needs ∼10 PSI to move the barriers and even less if moving one barrier at a time. Barrier air source will ideally be via the building’s air source. If unavailable, small commercial compressors can be used. However, these are loud during compression cycles and should be located outside the experimental rooms.

**Figure 4:**
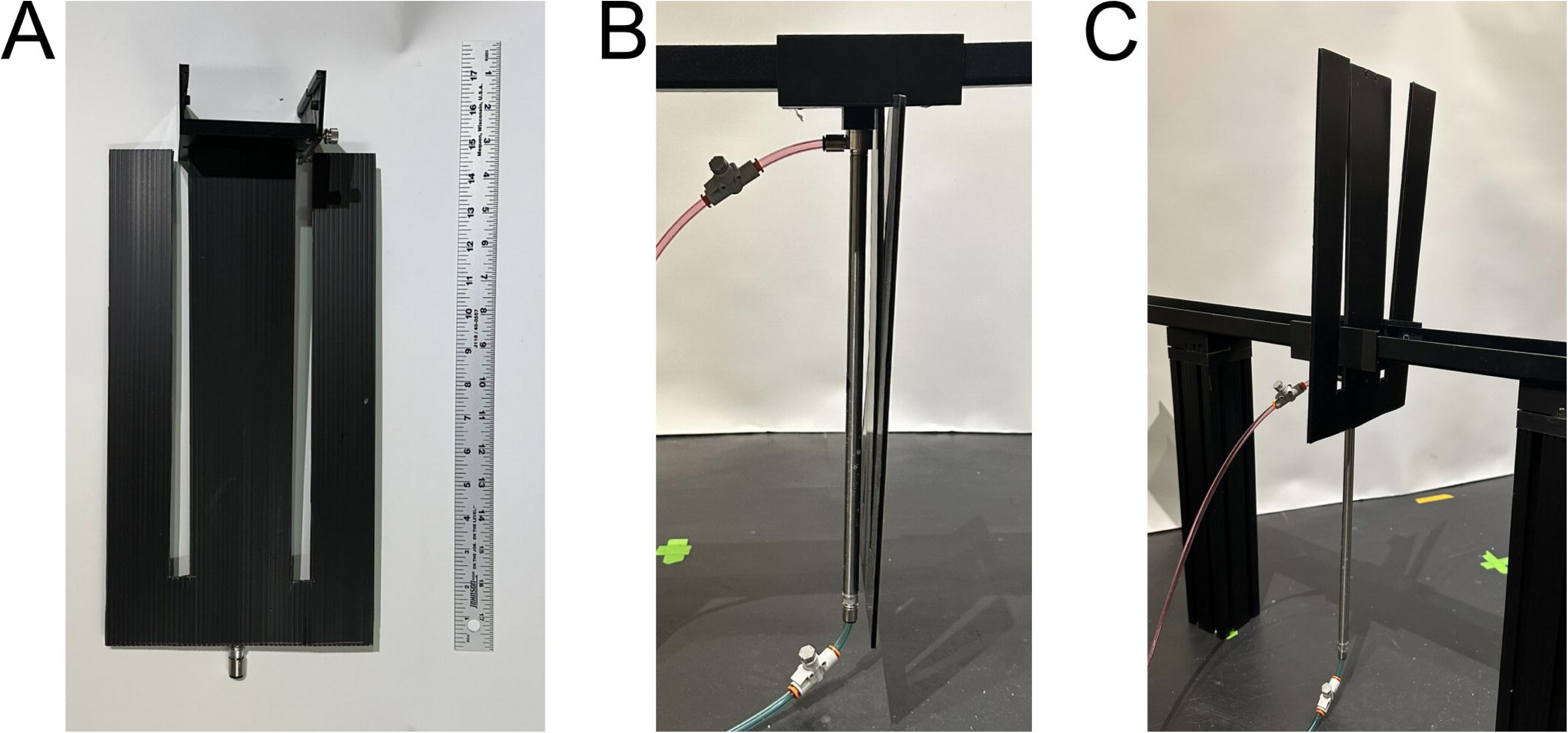
Automated Barriers. **A)** Front view of barrier assembly with corrugated plastic barrier attached to 3D printed track joint. **B)** Side view of barrier assembly in the down position between two track pieces. Flow regulators can be seen on tubing prior to the tubing’s connection to the piston. **C)** Barrier in the up position. **Figure 4-1** shows detailed part breakouts for barrier system including pneumatic control.

**Figure 4-1:**
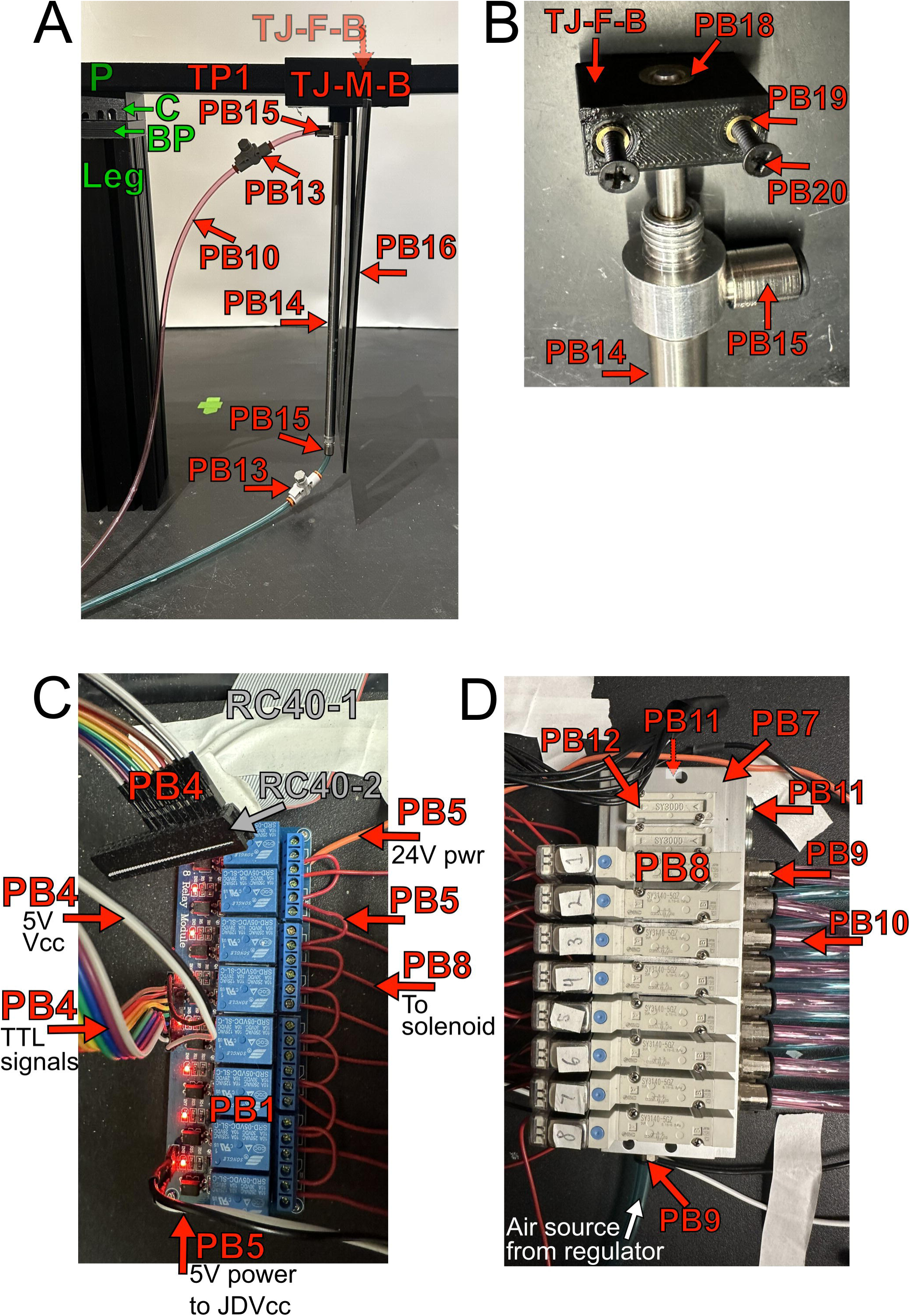
Automated Barrier parts. Part breakouts for **A)** barrier with track joint, **B)** barrier fastener, **C)** relay for barriers, and **D)** air manifold for Automated barrier system (Figure 4). Part IDs reference the parts list. For relay board (PB1) in **C**, do not use Vcc for power as it will pull power for the relays from your controller. Use an external power source (PB5) to JDVcc. Part ID colors correspond to the primary system they are associated with and are consistent with the Parts List’s tab colors. Red = track parts, green = reward wells, blue = reward delivery, grey = DIO connections. Low opacity indicates part is not visible in picture from this viewing angle.

#### Barrier relay circuit

We utilize relays so that the 5 V signals from the controller 1) can control the 24 V solenoids and 2) electrically isolate the solenoids from the controller. The circuit schematic can be seen in **Figure 2** and is pictured in **Figure 4-2**. This circuit daisy chains the 24 V power supplier to each relay’s common (COM) port meaning there will be two wires in each COM port except for the last one. You can then use either the normally open (NO) or normally closed (NC) ports to connect the positive lead to the solenoid. Depending on using the NO or NC, the barriers can start in the up or down position when the system is first powered on.

#### Maze assembly times

Initial setup of electronic parts can take a full week, such as soldering PCB components and hooking up connections from the hubs to the ECU / controller. However, once the hubs are hooked up in an experimental room, setting up usable mazes is quick. New users take ∼ 30 minutes to learn how to set up a linear track (3 straight tracks with small platforms at either end) for the first time. Assembly includes connecting the track pieces and legs, connecting the lick sensor and pump cables, and connecting the two syringes. Experienced users can assemble a linear track in approximately 10 minutes. To assemble an 8-arm radial maze with a barrier on each arm, it takes an experienced person an hour. The increased time is mainly due to all the tubing required for the barriers.

#### Cleaning procedures

Always consult your Institutional Care and Use Committee when developing cleaning protocols. Between each animal we remove feces and urine then clean the track pieces with bleach (10% sodium hypochlorite) followed by distilled water. Care is taken to ensure bleach does not get on the reward wells. Reward wells are cleaned of any excess reward with paper towels and water. At the end of each day, we remove the reward dispensing syringes from the maze and clean the tubing. First, the syringe plungers are pulled to remove excess reward. Second, a container is placed at the lower end of the reward well tube. We place the tip of a 60mL luer lock syringe full of water into the reward well hole. We designed the reward well hole is be sealed by the tip of the syringe. Approximately 20mL of water is then forced through the tubing and collected in the container. If viscous solutions such as high concentrations of sucrose or evaporated milk are used, we recommend cleaning the tubes with pipe cleaners once or twice a month to avoid build up. While rare with regular cleaning, it’s possible that the reward solution can build up on the IR emitter and/or receiver on the reward well. A stiff bristle brush and water can remove any build up.

### Experimental methods

All experimental methods were approved by the Brandeis University Institutional Care and Use Committee (#21001, #24001-A) or the Dartmouth College Institutional Care and Use Committee (#00002154). All experiments were conducted under the guidelines of the US National Institutes of Health. Data from one adult male TH-Cre rat (445g, 5 months old; Witten et al., 2011) is presented here, however, dozens of rats have carried out experiments on the AAM. The animal participated in a previous experiment (Ding et al., 2025) before participation in the radial arm maze associative memory task. Detailed methodological procedures including surgery are described in Ding et al., 2025. Briefly, the animal was singly housed due to neural implantation and given ad libitum access to food and water on a 12hr/12hr light/dark cycle. Lights were on from 7am – 7pm. Prior to training, the rat was food restricted to no less than 85% of his free feed weight. The rat was habituated to the experimenters and evaporated milk before being trained to shuttle back and forth on a linear track. Once readily shuttling, the rat underwent stereotaxic surgery and was injected with AAV5-EF1a-DIO-hChR2(E123T/T159C)-mCherry into the VTA and implanted with a custom hyperdrive consisting of tetrodes bilaterally targeting dorsal CA1 and prefrontal cortex and optrodes (optical fiber with tetrodes) to the VTA. The rat was given 3 weeks to recover while the virus expressed, tetrodes were lowered to their target areas, and he was retrained on the linear track. The rat was then trained on a W maze-based rule switching task.

After completion of the W maze rule switching paradigm, the rat was trained and tested on a sequence running task where each arm of the radial arm maze (**Figure 6A**) represented an item in the sequence. Utilizing the barriers, on a given trial the rat started at a Home arm and was presented two choice arms to choose from (**Figure 5**). Given the presented arms, the rat had to run the correct sequence from one to another. If correct, the rat received a reward at the first arm and the Home arm was blocked (*Barr 1*). The rat would then proceed to the second arm to get a second reward. After the second reward was delivered, the barrier to the first arm was raised (*Barr 2*) and the Home barrier was lowered (*Barr 1*) to start another trial.

**Figure 5:**
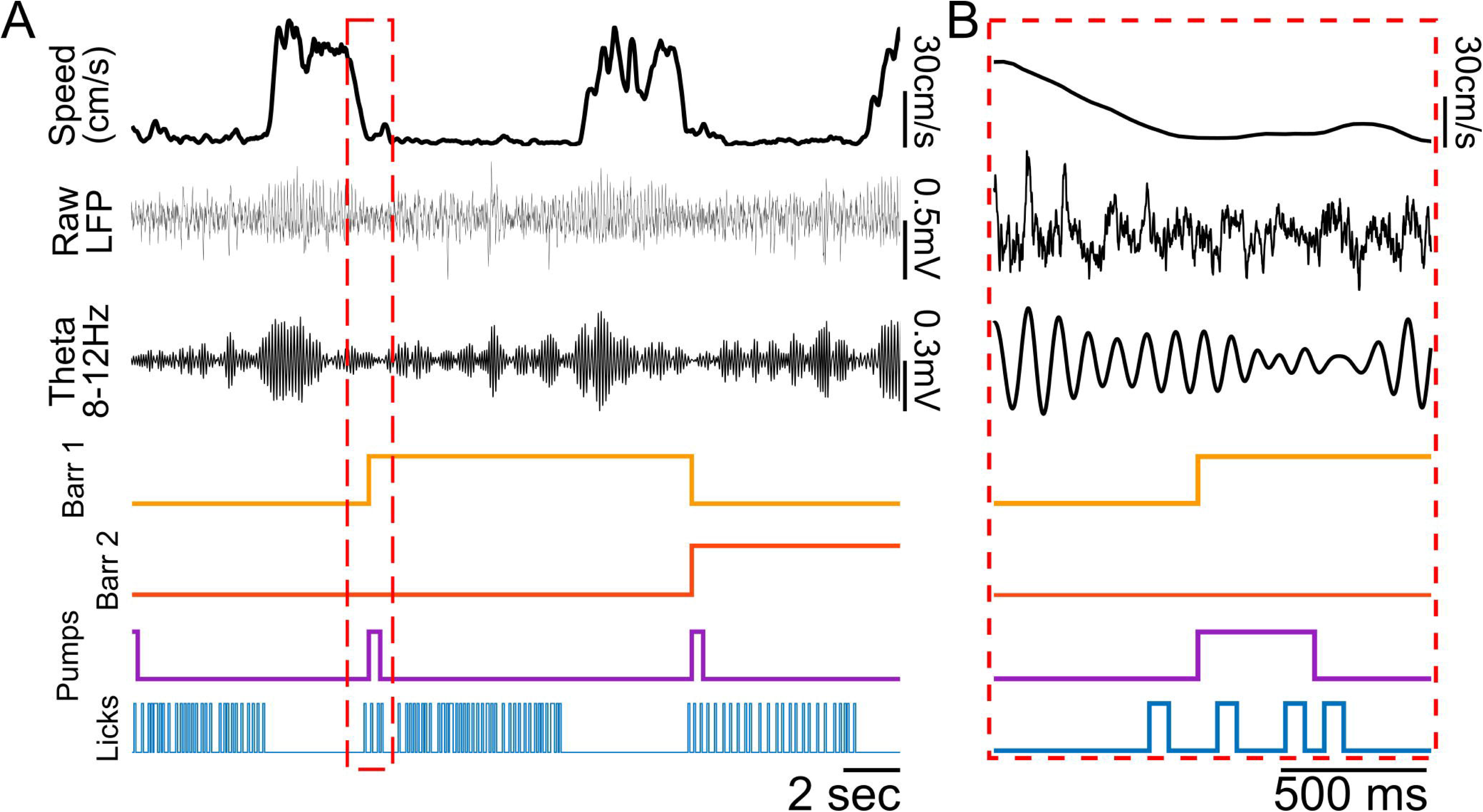
Example trial. **A)** Traces are from one trial of a rat running a memory sequence task on a radial arm maze. Top – Rat’s running speed as he consumed rewards and ran between arms. Raw LFP recorded from the hippocampus (dorsal CA1) and the LFP filtered for the theta rhythm (8-12 Hz)*. Barr 1 & 2* depict control signals for two different barriers as they are automatically controlled depending on the arm the animal is on. *High* signal indicates a raised barrier. Three reward pump signals were concatenated to one trace (*Pumps*, purple). A reward is dispensed while signals are high. The final trace shows putative licks detected from the integrated reward well IR beams at three different reward wells (*Licks*, blue). *High* indicates the beam was broken. **B)** Zoomed in portion of **A** (red dashed square) showing no artifact in the neural traces when both the barrier (yellow) and reward pump (purple) are turned on. Also depicted is our lick requirement where reward was only dispensed once a rat initiated licking to indicate their choice.

**Figure 6:**
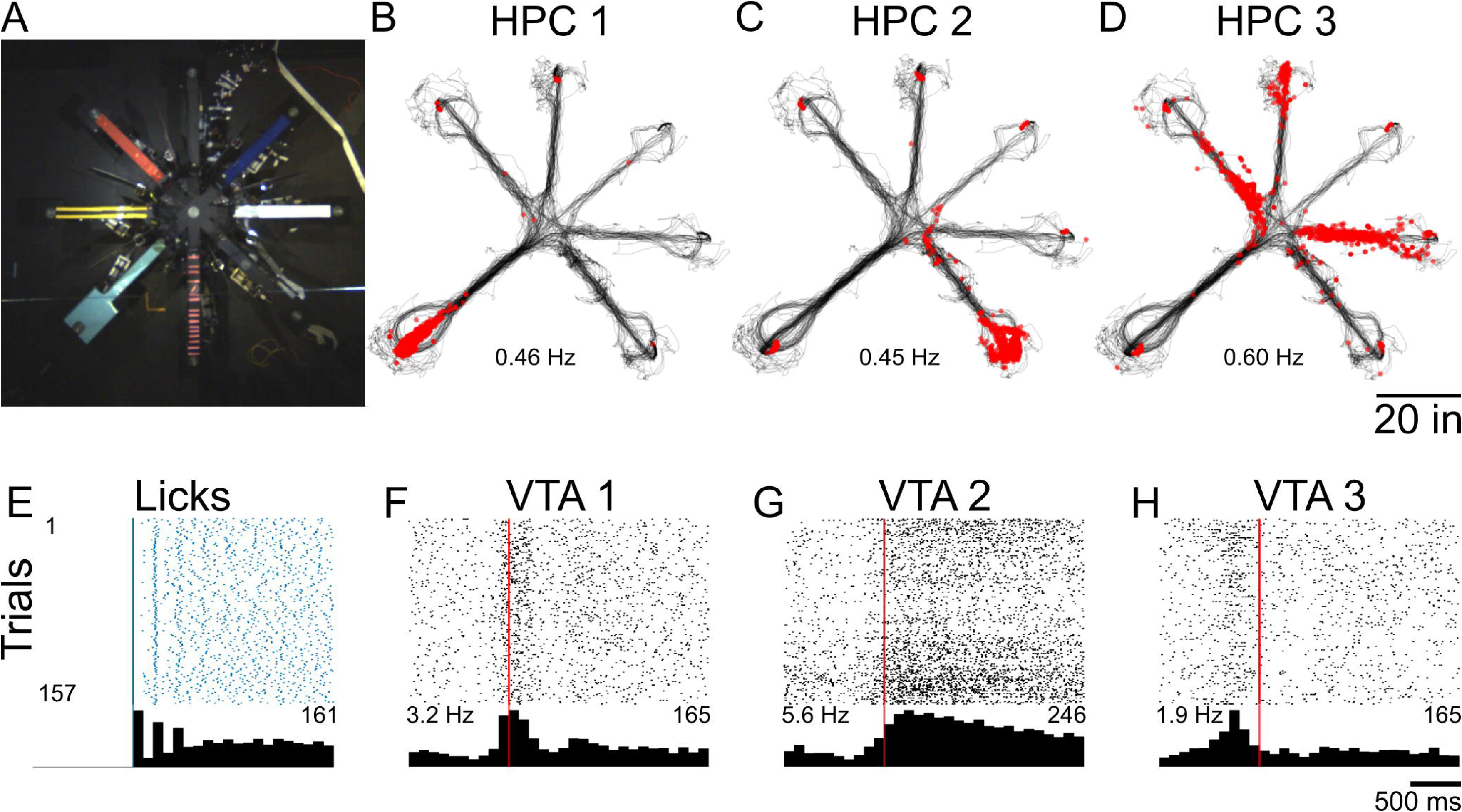
Example neuronal activity on an AAM task. **A)** 8-arm radial arm maze used to test sequence memory in rats. **B-D)** Example plots of three HPC neurons from one recording session on the radial arm maze. Black lines depict the rat’s position on the maze. Each red dot is the location where an action potential was fired. Mean firing rates for the recording sessions shown below each plot. Neurons shown in **B** and **C** are spatially selective, firing to the end platforms of an arm. The cell in **D** shows selective spatial firing to multiple arms. **E)** Raster plot and peri-event time histogram (PETH) centered on the first lick of correct trials from the same recording session across all reward wells. Inset right is peak lick count. **F-H)** Rasters and PETHs of three example VTA neurons around reward consumption. Red vertical line indicates first lick. Mean firing rates for the recording session are inset on the left while max spike count on the histogram is inset right. A variety of responses were observed including phasic response at reward delivery (**F**), tonic response during licking (**G**), and preemptive firing before lick onset (**H**).

The session was recorded with an overhead camera (Mako G-158C, Allied Vision Inc.) recording at 30 frames per second. The speed of the rat was calculated by tracking an LED on the headstage (SpikeGadgets) of the animal in DeepLabCut (Nath et al., 2019) and calculating its instantaneous velocity and smoothing with a centered moving average of 7 frames. Hippocampal local field potential data was acquired via a SpikeGadgets MCU and exported with a lowpass filter with a cut off of 500 Hz. Theta was filtered at 8 – 12Hz using a third order Butterworth filter. Spiking data was obtained using MountainSort (Chung et al., 2017) and manually curated with Plexon OfflineSorter (Plexon Inc.) primarily using principal components and peak-valley distance to visually inspect cluster quality.

## Results

We designed a maze system using modular track pieces (**Figure 1A**) that enables the creation of a wide array of two-dimensional track environments (**Figure 1C; See Figure 1-1 for complex mazes**). All the necessary files and materials can be found on the Gitlab repository (https://gitlab.com/jadhavlab/AdaptAMaze/-/tree/main?ref_type=heads). We chose to make the track pieces out of aluminum for a variety of reasons. Using metal ensures the track pieces are strong, long-lasting, non-porous, easy to clean, and minimize static buildup on the animals. We have noted during low-humidity periods that static discharge can occur. Increasing the room’s humidity ameliorates the issue. We specifically selected aluminum as it is an inexpensive sheet metal that is lightweight and corrosion resistant (comparative to stainless steel). It’s a soft metal that allows for modifications if needed with simple tools (e.g. handheld drill or hacksaw). Furthermore, aluminum can be anodized to a matte black surface, as we have done, to minimize light reflections to improve animal tracking. Each track piece is supported by a leg, and track pieces are joined using custom 3D-printed track joints (**Figure 1B**). Track pieces are designed to meet at locations on an 18-inch grid (except for the platform pieces), thereby allowing any combination of pieces to be used to create the maze of choice. The system is scalable, as mazes can be expanded by simply adding another piece. It is also flexible, as a maze can be adapted by replacing one or more pieces by simply disconnecting the joint connections and placing alternative track pieces. Due to the standardized design and componentry, maze environments can be easily recreated both in and across labs, greatly enhancing repeatability.

All aspects of the AAM system can be automated including lick detection, reward dispensing, and barrier movements (**Figure 2, 2-1**). The maze system features automated reward well lick detection and liquid reward distribution (**Figures 3, 3-1, and 3-2**). A custom printed circuit board (PCB; **Figure 3A**) routes 5V signals from detected licks from each reward well to the digital input/output (DIO) controller (e.g. Environmental Control Unit or “ECU” from SpikeGadgets, Arduino, Raspberry pi, etc.). Integrated into each reward well is an infrared beam break circuit, two visible signaling LEDs, and tubing for liquid reward delivery (**Figure 3B, C**). Each track piece is held in place by either a reward well or a flat “plug” that does not have a well or hole (not shown). A reward well or plug is screwed into the leg connector (**Figure 3D**) attached to the supporting leg baseplate (**Figure 3E**). These components are tucked into the leg assembly upon construction (**Figure 3E,F**), protecting componentry from curious rodents and allowing for quick connections and easy cleaning. Liquid rewards were chosen to minimize chewing artifacts common to solid food rewards (Fabietti et al., 2020; Mondragón-González & Burguière, 2017). Custom software controls the SpikeGadgets ECU for reward programs. Further control over reward delivery can be programmed to avoid. For example, we program a lick requirement so that reward is dispensed only after a rat licks a certain number of times at a well. This requirement serves as a behavioral readout of the rats choice and prevents unwanted events from dispensing milk, such as the rat standing on the well. A similar PCB controls the visible LEDs in the reward wells and a third PCB controls the pumps to deliver liquid rewards (all PCBs, including circuit diagrams: https://gitlab.com/jadhavlab/AdaptAMaze/-/tree/main?ref_type=heads). If desired, an Arduino can interface directly to the PCBs to log licks or deliver rewards.

To achieve environmental flexibility within a recording session, the maze system includes the ability to place automated barriers between any two track pieces (**Figures 4, 4-1**). Barriers are pneumatically driven to avoid electrical noise commonly generated from electric motors. Like the reward system, barrier position is controlled through digital inputs from the ECU to a simple relay interfacing with solenoids to direct airflow to the appropriate barrier. Barrier speed can be individually and directionally (i.e., up and down) controlled via flow regulators (**Figure 4 B,C**). Barriers can be made to a variety of heights but are limited by piston length (McMaster-Carr supplies these pistons with stroke lengths up to 36 in) and maze leg height. Commonly, barriers are constructed from corrugated plastic which is easy to work with, inexpensive, and cleanable. Clear Plexiglas barriers have been used to block routes while maintaining visual cues. However, any material could be used as a barrier with sufficient air pressure. Overall, due to the modular design of the AAM, certain components such as the reward wells or lick sensors could be adapted by other labs for their mazes (e.g. the automated barriers were used in two different labs, see **Movie 1**).

The AAM has been extensively tested over the past five years in our lab. We have used the AAM in a variety of configurations including linear tracks, W mazes, 8-arm radial mazes, and other complex configurations (**Figure 1-1**). The mazes have been tested exclusively with freely behaving rats (male and female Long Evans aged 3 to 13 months). Researchers utilizing mice may want to modify the track pieces to be smaller. Some rats were implanted with hyperdrives, optical fibers, or both to a variety of brain regions including the hippocampus (HPC), prefrontal cortex, and ventral tegmental area (VTA). **Figure 5** depicts example data from one trial from one animal running a radial arm maze (**Figure 6A**) that had automated barriers to control arm access (see **Movie 2** for example trial). We have utilized the AAM to investigate a wide range of system neuroscience questions including the hippocampal representations of complex sequences (**Figure 6B-D**) and VTA neuronal responses around reward delivery over learning (**Figure 6E-H**). We look forward to the limitless possibilities other researchers come up with using and adapting the AAM system.

## Discussion

Here we have detailed a novel modular maze system for behavior and systems neuroscience research in rodents. The design integrates reward ports into the track pieces and includes automated lick detection and liquid reward distribution. Automated barriers can be included between any two track pieces, furthering flexibility of the design. Together, our open-source design comprises a simplified system capable of replicating standard maze configurations as well as novel experimental designs.

Notably, this design allows for the construction of stereotyped and clearly-documented maze designs within and across labs, increasing repeatability within an experiment and the ability for labs to replicate the same experimental setups. For example, classic spatial navigation and working memory assays on environments such as ‘T’, ‘Plus’, and ‘W’ mazes can be constructed with this system to exact specifications and with shared, standardized control code. These same setups can then be exactly replicated without variability in implementation due to varied hardware (e.g. material, port type) or behavioral task control code. Furthermore, the automation of reward delivery and barriers could potentially reduce variability in experiments that previously had experimenters manually releasing rewards or adjusting barriers.

Another benefit of a modular system is the flexibility to reuse lab spaces and adapt maze designs with ease. A common maze setup is a single-piece or fixed design. For another researcher to then use that same experimental space with a different maze design, the first maze must be relocated. Obviously, this limits the feasible number of actual mazes used in a lab. In addition, if two or more mazes want to be used for the same experiment session, it may be difficult to make the transition in a timely manner. With our system, this can be addressed quickly and easily with the repositioning or exchanging of track pieces, facilitating both minor space manipulation experiments in minutes or complete track redesigns in tens of minutes for a single experimenter. The modular aspect also facilitates simple scaling of mazes by adding segments to existing track pieces, something impossible with fixed or single-piece designs.

Perhaps the most exciting benefit of this system is the reduced financial, engineering, and setup costs of experiments. Spurring the AAM was the inordinate costs many companies charge for prebuilt automated mazes that range from thousands to tens of thousands of dollars and often require additional integrations (e.g. lick sensors) at additional cost. Furthermore, many commercial mazes use proprietary software that can limit integration into other systems (e.g. electrophysiology rigs) and impede reproducibility. In March 2025, we were quoted $27,744 USD for an automated 8-arm rat maze from a well-known maze company. In contrast, an AAM system for an 8-arm radial maze purchasing 10 pieces of straights (I) and platforms (Plat_small) and two 8-arm intersections (Oct) would cost approximately $9,500 (price assumes the lab has 3D printers). Specifically, the track pieces would cost approximately $2,000, legs $750, barriers $1,800, reward delivery $4,300, and smaller parts making up the remainder (see Parts List for detailed pricing). Substantial savings (∼ $4,000) are possible by using peristaltic pumps ($200) rather than syringe pumps. Importantly, the same track pieces could be reused to construct multiple linear tracks, two 4-arm plus mazes, and three T/Y mazes. The AAM system also reduces new maze prototyping times by months or even years. Our open-source designs mean researchers could adapt our existing track pieces, reward wells, barriers, and circuits to meet their needs. For example, the lick sensors could be adapted to nose poke sensors and the barrier system could be made into an air puff system. Because of the minimal cost of any particular configuration, many configurations can be quickly piloted to find the preferred environmental design.

This maze system is compatible with behavioral recording as well as neurophysiology recording techniques, including *in vivo* electrophysiology and one-photon calcium imaging. System components are entirely below the track surface, clearing the space above the track for unimpeded visual recording or physical tethering. In particular, this design will enable continuous neural monitoring during behavioral tasks in multiple maze environments.

The Adapt-A-Maze does however have some limitations. Simple translations or rotations of the maze in the room are difficult with the current design, as the connections between track segments are weak compared to the weight of the legs. One possible solution is to connect the legs with additional T-slotted beams to add strength, but this will trade off with the amount of time to adapt the system. Two-dimensional mazes are very stable on their own due to the weight of the legs and the track pieces bracing one another. However, linear tracks can be unstable in the sense that a forceful bump could knock it over. In tight quarters where the risk of bumping a linear track may be high, using T-slot accessories to widen the leg base may be helpful. Another concern is the extent of design flexibility. Our maze system is currently laid out on a grid system for ultimate flexibility. As is, the grid does limit the potential maze designs for certain radial configurations. However, users can easily modify or create new track pieces or customize components to add functionality to the system. The only compatibility requirements are to interface with either our track joint and reward well designs or the basic sheet metal track pieces. Finally, the current track pieces are incapable of three-dimensional maze designs, so unique mazes (Grieves et al., 2020; Wilson et al., 2015) will still be required to explore three-dimensional navigation. Three-dimensional *surfaces* could be possible with varied leg heights and bends in the track pieces.

We have encountered some common wear and tear items that may be useful to point out. Some of our mazes utilize the barriers every trial, resulting in hundreds of up-down cycles per day. Over time, the barrier material (if using corrugated plastic) can dislodge from the air cylinder. Daily checks to examine their stability are helpful. Even so, replacements are only needed every year or so. Secondly, a subset of rats may become frustrated when not receiving a reward at a well and will attempt to chew it. We considered this in the reward well design by utilizing smooth surfaces. However, given enough time, their teeth can scratch the plastic enough it’s worth replacing to ensure cleanability.

There are many alternative possibilities with different relative costs and benefits. Single-piece mazes made from materials such as wood are quick and cheap to construct but are permanent in their design and occupy considerable experimental space when not in use. Further, some institutions do not allow wooden mazes for animal use due to being porous and hard to clean. Complex, unique mazes can be perfectly catered to a single experiment, but often have immense engineering setup costs and permanent designs. Another maze alternative is a virtual reality (VR) setup. The utility of VR setups as compared to our design is similar to other real world mazes (Chen et al., 2018; Minderer et al., 2016). VR offers the ability to precisely control sensory variables and even decouple typically-coupled variables such as movement speed and visual flow, but that same decoupling can also be a disadvantage and creates an unnatural situation that may not be desired for the study. VR setups can also be more amenable to head-fixed recording techniques, though simple mazes are now possible on head-fixed setups (Kislin et al., 2014). Finally, an impressive modular track system was recently published by Hoshino and colleagues (Hoshino et al., 2020). Their system boasts many of the features of our system, but with a few important distinctions. Their reward system is for solid food as opposed to liquid, and their barrier system is not automated. Hoshino et al. also use a grid system but it is implemented on the floor, limiting their ability for configurations to 45– and 90-degree segments. They have developed a beam break system that is decoupled from reward locations, allowing for more flexible behavioral control than currently available with our AAM system. Many of these variations could be added to our system due to its fundamental simplicity, but they are not currently integrated.

Mazes remain a ubiquitous tool in rodent behavior and systems neuroscience (Wijnen et al., 2024). We have designed a modular maze system to enable flexible maze designs and rapid experiment prototyping and development. Adaptation of this standardized system will also allow for improvements in repeatability and replication. Overall, the authors believe this maze system adds an important tool for researchers and will facilitate and expedite novel behavior and systems neuroscience discoveries.

## Author Contributions

All authors contributed to the design of the AAM system. B.S.P., J.M.O., C.A.L., E.D., and J.H.B. performed the research of prototyping the AAM. B.S.P. analyzed the data. B.S.P. and J.M.O. wrote the manuscript and all authors edited it.

## Acknowledgements

We thank the Brandeis NSF I-Corps Program for guidance during product development, our I-Corps teammates Xin Yao Lin and Faye Raymond for helpful discussions during development, Francisco Mello for his assistance in fabrication of maze components, Andrew Alvarenga (Grasshopper Machine Werks, LLC.) for technical assistance on the pneumatic barriers. We thank many of our colleagues who helped to develop the maze, especially the barriers, including Dwayne “Whitey” Adams, Shahaf Weiss, Emily Aery Jones, Ryan Young, Caine Rees, Noah Moore, Jianing Yu, and Kevin Allen. We would also like to thank all members of the Jadhav and van der Meer labs for helpful comments and feedback during the design and implementation of the maze system.

## Movie legends

**Movie 1:** Example of implanted rat behavior using the automated barrier system in the van der Meer lab. A male Long-Evans rat was trained to shuttle between the two ends of the linear track for food; later on, a delay zone was added that required the rat to nosepoke for a certain duration in order for the barrier to lower. In this case, the rat was implanted with a 32 tetrode hyperdrive connected to a SpikeGadgets recording system, which can be used seamlessly with the barriers.

**Movie 2:** Example trial of rat running on an AAM 8-arm radial maze, similar to those shown in Figures 5 and **6**, in the Jadhav lab. Rat was implanted with a 64 tetrode drive and tethered while running. Neural data was acquired with a SpikeGadgets system that also controlled the automation of the maze. The rat had no issues with the barrier sounds nor the tether getting caught on any part of the maze.

